# A single dose of dietary nitrate increases maximal muscle speed and power in healthy older men and women

**DOI:** 10.1101/564849

**Authors:** Andrew R. Coggan, Richard L. Hoffman, Derrick A. Gray, Ranjani N. Moorthi, Deepak P. Thomas, Joshua L. Leibowitz, Dakkota Thies, Linda R. Peterson

**Author notes:** Address correspondence to: Andrew R. Coggan, PhD, Department of Kinesiology, Indiana University Purdue University Indianapolis, IF 101C, 250 University Boulevard, Indianapolis, IN 46112.

## Abstract

**Background:** Aging results in reductions in maximal muscular strength, speed, and power, which often lead to functional limitations highly predictive of disability, institutionalization, and mortality in the elderly. This may be partially due to reduced nitric oxide (NO) bioavailability. We therefore hypothesized that dietary nitrate (NO_3_^−^), a source of NO via the NO_3_^−^ → nitrite (NO_2_^−^) → NO enterosalivary pathway, could increase muscle contractile function in older subjects.

**Methods:** Twelve healthy older (age 71±1 y) subjects were studied using a randomized, doubleblind, placebo-controlled, crossover design. After fasting overnight, subjects were tested 2 h after ingesting beetroot juice containing or devoid of 13.4±0.5 mmol NO_3_^−^. Plasma NO_3_^−^ and NO_2_^−^ and breath NO were measured periodically, and muscle function was determined using isokinetic dynamometry.

**Results:** NO_3_^−^ ingestion increased (P<0.001) plasma NO_3_^−^, plasma NO_2_^−^, and breath NO by 1050%, 140%, and 110%, respectively. Maximal velocity of knee extension increased (P<0.01) by 1.1±0.3 rad/s, or by 10.9±3.5%. Maximal knee extensor power increased (P<0.05) by 0.14±0.06 W/kg, or by 4.4±2.3%.

**Conclusions:** Acute dietary NO_3_^−^ intake improves muscle contractility in older individuals. These findings have important implications for this population, in whom diminished muscle function can lead to functional limitations, dependence, and even premature death.

---

Aging is accompanied by a progressive reduction in the maximal strength, speed, and especially power of skeletal muscle. These age-related physiological changes often lead to functional limitations that are highly predictive of disability, institutionalization, and mortality in the elderly (1,2). Thus, any intervention that significantly enhances muscle contractile function could potentially improve the health, quality of life, and possibly even the longevity of older individuals.

Numerous factors undoubtedly account for the decline in muscle contractility described above, with age-related changes in the size, properties, and neural control of muscle likely all playing a role (3). Another important factor, however, may be a reduction in nitric oxide (NO) bioavailability with aging. Although initially identified as a vasodilator, i.e., as “endothelium-derived relaxing factor”, NO is in fact a key cellular signaling molecule with pleiotropic effects in many tissues. These include skeletal muscle, wherein among other effects NO significantly increases the rate of force development, maximal shortening velocity, and maximal power during concentric contractions (4,5). With aging, however, whole-body NO production decreases, as evidenced, e.g., by a progressive decline in the plasma concentrations of its downstream metabolites, nitrite (NO_2_^−^) and nitrate (NO_3_^−^) (6,7). In skeletal muscle itself, there is a dramatic decrease in interstitial NO_2_^−^ and NO_3_^−^ concentrations (8), as well as a decline in the activity of the neuronal form of NO synthase (NOS) (9,10), the primary isoenzyme within muscle responsible for the synthesis of NO from L-arginine, O_2_, and NADPH. These changes are accompanied by an age-related reduction in flow-mediated vasodilation (11,12), perhaps the hallmark indicator of NO bioavailability. In turn, the latter (i.e., flow-mediated vasodilation) has been shown to correlate positively with muscular power and physical functioning in older men and women (13). Taken together, these data suggest that decreased NO production may contribute to the age-associated decline in muscle contractile properties and hence in functional capacity.

Along with the NOS pathway, NO can also be produced in the body via chemical reduction of NO_3_^−^ to NO_2_^−^ and hence to NO (14). This NO_3_^−^ → NO_2_^−^ → NO pathway is facilitated by commensal bacterial nitrate reductases in the mouth and/or endogenous mammalian nitrate reductases (e.g., xanthine oxidase) in the tissues, and unlike the NOS pathway is accelerated at low pO_2_ and low pH, conditions typical of resting and especially contracting muscle (15). Indeed, muscle NO_3_^−^ has recently been shown to be an important source of whole-body NO production (16). In turn, muscle NO_3_^−^ can be replenished via oxidation of NO to NO_2_^−^ to NO_3_^−^, or obtained via the diet (17). Numerous studies in recent years have therefore examined the effects of dietary NO_3_^−^ supplementation on various physiological responses mediated by NO, e.g., muscle blood flow (18). This includes studies of the effects of NO_3_^−^ ingestion on the responses to *aerobic* exercise in older individuals (19,20). To date, however, no study has determined the influence of dietary NO_3_^−^ on muscle contractile function in older humans.

We have previously demonstrated that NO_3_^−^ supplementation increases muscle speed and power in healthy younger and middle-aged men and women (21,22), athletes (23), and patients with heart failure (HF) (24). However, it cannot simply be assumed that dietary NO_3_^−^ will be equally efficacious in older individuals. For example, in rodents the effects of dietary NO_3_^−^ are more prominent in fast-vs. slow-twitch muscle (4,5), and some (25,26), but not all (27), studies of humans have found that aging results in a reduction in the percentage of fast-twitch muscle fibers. Alternatively and/or in addition, an age-related decrease in the ability of salivary glands to concentrate NO_3_^−^ (28) or more rapid destruction of NO due to increased production of reactive oxygen species in aged muscle (29,30) could limit the beneficial effects of dietary NO_3_^−^ in older persons.

The purpose of the present study was to test the hypothesis that by enhancing NO production, acute dietary NO_3_^−^ supplementation would improve muscle contractile properties in healthy older men and women. If so, this would have important implications for this population, in whom diminished muscle function can contribute significantly to functional limitations, disability, dependence, and even death (1,2).

## Methods

### Subjects and experimental design

Twelve men and women (n=6 each) with mean (±SE) age, height, weight, and body mass index of 71±1 y, 1.71±0.03 m, 75.2±3.2 kg, and 25.7±1.0 kg/m^2^, respectively, were studied using a double-blind, placebo-controlled, crossover design. The first six were studied in St. Louis, with the remaining six studied in Indianapolis using essentially identical procedures. All were in good health, as assessed by physical exam and standard blood chemistries, including fasting glucose and insulin levels and blood lipid profile. Exclusion criteria included age <65 or >79 y or history of significant cardiovascular disease (e.g., stage II or greater hypertension, heart failure, myocardial infarction/ischemia, myocardial/pericardial diseases, moderate or severe valvular disease), renal disease (eGFR <60 mL/min/1.73 m^2^ or 61-90 mL/min/1.73 m^2^ and albumin:creatine ratio >30), liver disease (SGOT/SGPT >2x normal), anemia (hematocrit <30%), or any other contradiction to strenuous exercise. Those taking phosphodiesterase inhibitors (e.g., sildenafil) were also excluded, as these can potentiate NO effects (31). Finally, those taking proton pump inhibitors, antacids, xanthine oxidase inhibitors, or on hormone replacement therapy were also excluded, as these can affect reduction of NO_3_^−^ and NO_2_^−^ (32,33). Approval for the study was obtained from the Human Research Protection Office at Washington University in St. Louis and the Human Subjects Office at Indiana University, and written, informed consent was obtained from each subject.

### Experimental protocol

Subjects reported to the Clinical Research Unit at Washington University in St. Louis or the Clinical Research Center at Indiana University Purdue University Indianapolis in the morning after avoiding high NO_3_^−^ foods (e.g., arugula, beets, spinach) for the previous 24 h and any food, caffeine, and alcohol for the previous 12 h. During this period they also refrained from chewing gum, brushing their teeth, or using an antibacterial mouthwash, as these can block reduction of NO_3_^−^ to NO_2_^−^ by oral bacteria (34). After measurement of seated heart rate and blood pressure and the level of NO in breath (NIOX VERO^®^, Circassia Pharmaceuticals, Mooresville, NC), an intravenous catheter was placed and a baseline blood sample was obtained for subsequent measurement of plasma NO_3_^−^ and NO_2_^−^ concentration using high performance liquid chromatography (ENO-30^®^, Eicom USA, San Diego, CA). The subject then ingested either 1) a commercial beetroot juice (BRJ) supplement (Beet It Sport^®^, James White Drinks, Ipswich, UK) containing (based on direct measurement) 13.4±0.5 mmol of NO_3_^−^, or 2) an equal volume of BRJ from which the NO_3_^−^ had been removed by the manufacturer. The subject then rested quietly, with additional measurements of heart rate, blood pressure, breath NO, and plasma NO_3_^−^ and NO_2_^−^ obtained 1 and 2 h after BRJ ingestion. Muscle contractile function was then measured during the 3^rd^ hour post-BRJ ingestion as described below, with final measurements of hemodynamics and NO3VNO2VNO obtained 10 min post-exercise. The subjects were then fed a light meal and released from the CRU/CRC. After a 1-2 wk washout period, they returned for second visit during which they were “crossed over” to the other treatment and all of the above procedures repeated.

### Measurement of muscle contractile function

The maximal power (Pmax) and velocity (Vmax) of the quadriceps muscle group were measured using an isokinetic dynamometer (Biodex System 4 Pro^®^, Biodex Medical Systems, Shirley, NY) as previously described in detail (21). Briefly, after alignment of the lateral femoral epicondyle of the subject’s dominant leg with the axis of rotation of the dynamometer, they performed maximal knee extensions at angular velocities spanning the ascending limb of the power-velocity relationship, i.e., at 0, 1.57, 3.14, 4.71, and 6.28 rad/s (0, 90, 180, 270, and 360 °/s). Subjects executed 3-4 maximal efforts at each velocity and were allowed 2 min of rest between each set of contractions. Isometric testing was conducted at a knee joint angle of 1.22 rad (70°), thus approximating the angle at which peak torque was achieved during the isokinetic contractions. Peak power at each velocity was calculated by multiplying peak torque by velocity, after which Pmax and Vmax were determined by fitting an inverted parabola to the peak power-velocity relationship (21). After the final 2 min rest period, subjects performed 50 maximal knee extensions at 3.14 rad/s (total test duration approximately 1 min) to determine the effects of dietary NO_3_^−^ on muscle fatigue resistance during intense muscular contractions in older individuals. Subjects were instructed to exert themselves maximally during each of the 50 knee extensions and vociferously encouraged to do so throughout the test.

### Statistical analyses

Normality of data distribution was tested using the D’Agostino-Pearson omnibus test. Hemodynamic, plasma NO_3_^−^ and NO_2_^−^, breath NO, peak torque, and peak power data from the placebo and NO_3_^−^ trials were compared using two-way (treatment × time or treatment × velocity) repeated measures ANOVA. Post-hoc testing was performed using the Šidák multiple comparison procedure. Absolute values and changes in Vmax and Pmax were compared between treatments using two-sample paired and one-sample t tests, respectively. Effect sizes were calculated as partial eta squared (ηp). P<0.05 was considered significant. All statistical analyses were performed using GraphPad Prism version 8.01 (GraphPad Software, La Jolla, CA).

## Results

All subjects completed the entire study protocol. One individual experienced mild-to-moderate diarrhea during both the placebo and NO_3_^−^ trials, but no other adverse effects were observed.

Values for plasma NO_3_^−^ and NO_2_^−^ and breath NO are shown in Table 1. There were significant main effects for time 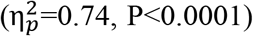 and treatment 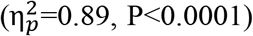 and a significant time × treatment interaction effect 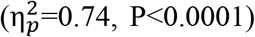 for plasma NO_3_^−^. Post-hoc testing indicated that plasma NO_3_^−^ was higher (P<0.0001) in the NO_3_^−^ trial at 1 and 2 h following BRJ ingestion and also at 10 min post-exercise. Similar results were observed for plasma NO_2_^−^, with time 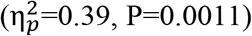, treatment 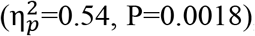, and time × treatment interaction effects 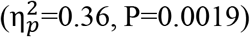 all being significant. Plasma NO_2_^−^ was significantly different between trials at 2 h (P=0.016) and 10 min post-exercise (P<0.0001). Finally, for breath NO there were significant main and interaction effects (time: 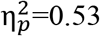, P=0.0003; treatment: 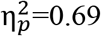, P=0.0013; interaction: 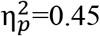, P=0.0004), with breath NO being higher in the NO_3_^−^ vs. the placebo trial at 1 and 2 h post-ingestion and 10 min post-exercise (P=0.0002, P=0.0009, and P<0.0001, respectively).

**Table 1.** Effects of dietary NO_3_^−^ on plasma NO_3_^−^, NO_2_^−^, and breath NO in healthy 65-79 y old men and women.

|  |  | Time of measurement |  |  |  |
| --- | --- | --- | --- | --- | --- |
|  | <u>Trial</u> | Pre | 1 h post | 2 h post | 10 min post |
|  |  | <u>ingestion</u> | <u>ingestion</u> | <u>ingestion</u> | <u>exercise</u> |
| Plasma NO<sub>3</sub><sup>-</sup><br>(μmol/L) | Placebo | 47 | 40 | 45 | 44 |
|  |  | ±9 | ±6 | ±8 | ±8 |
|  | Nitrate | 39 | 422‡ | 453‡ | 446‡ |
|  |  | ±5 | ±56 | ±62 | ±58 |
| Plasma NO<sub>2</sub><sup>-</sup><br>(μmol/L) | Placebo | 0.27 | 0.28 | 0.31 | 0.28 |
|  |  | ±0.04 | ±0.03 | ±0.04 | ±0.04 |
|  | Nitrate | 0.31 | 0.44 | 0.52* | 0.72‡ |
|  |  | ±0.04 | ±0.08 | ±0.06 | ±0.15 |
| Breath NO<br>(ppb) | Placebo | 21 | 24 | 26 | 22 |
|  |  | ±3 | ±3 | ±3 | ±3 |
|  | Nitrate | 22 | 41† | 41† | 49‡ |
|  |  | ±3 | ±5 | ±5 | ±8 |

Note. Values are mean±SE for n=12.
Nitrate trial significantly higher than placebo trial: \*P<0.05, †P<0.001, ‡P<0.0001.

Heart rate and blood pressure data are provided in Table 2. For heart rate, there was a significant main effect for time 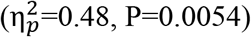, but not for treatment 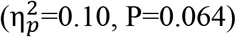 or for time × treatment interaction 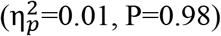. Systolic blood pressure also differed over time 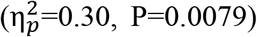, but the treatment 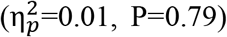 and interaction 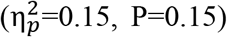 effects were non-significant. Diastolic blood pressure, on the other hand, did not vary with time 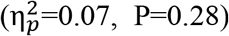, but did vary with treatment 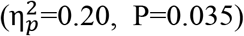. The time × treatment interaction effect was not significant 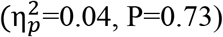.

**Table 2.**
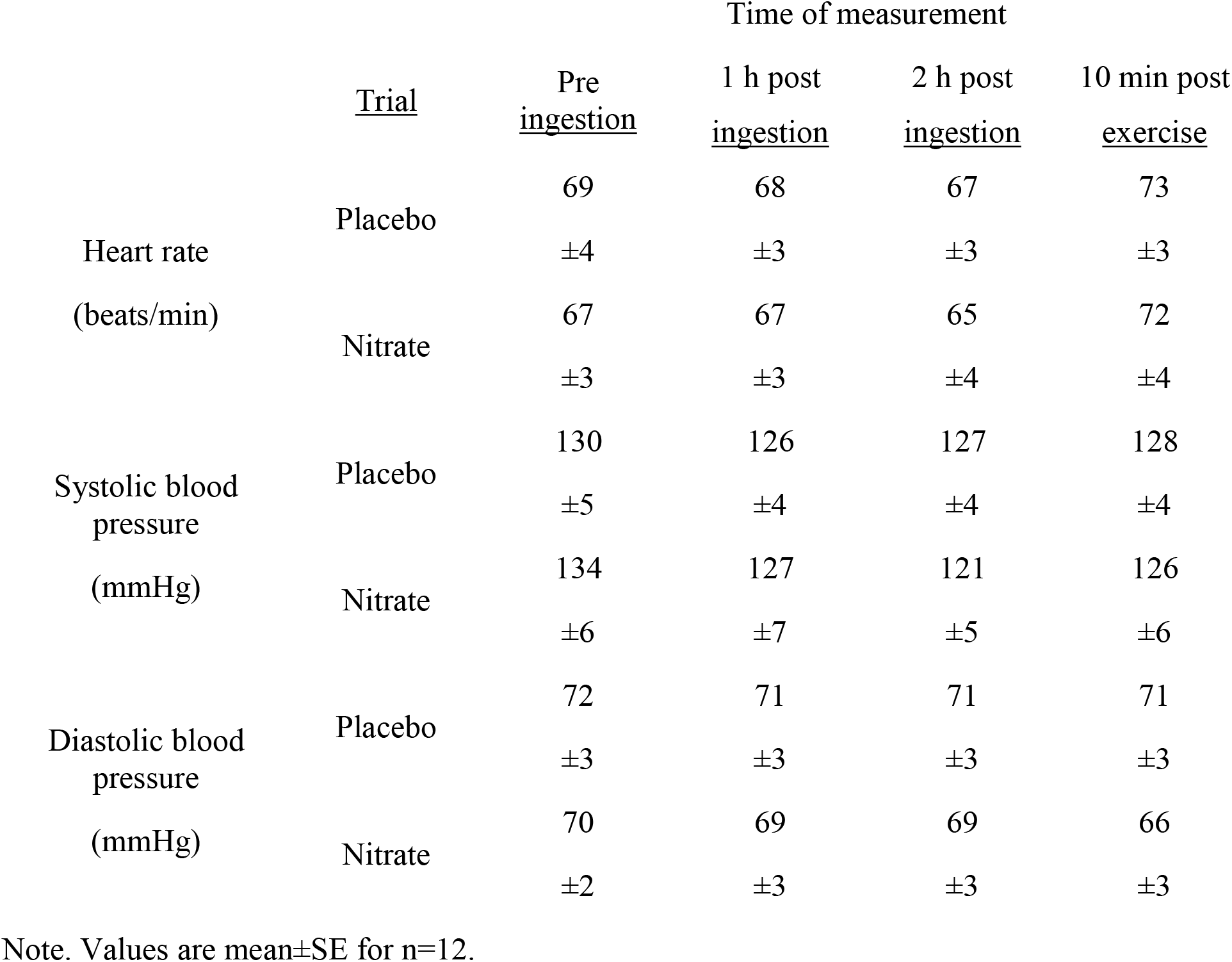
Effects of dietary NO_3_^−^ on heart rate and blood pressure in healthy 65-79 y old men and women.

Data for peak knee extensor torque and power are shown in Table 3. For absolute peak torque, there was a significant effect for velocity 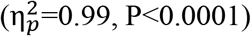, a non-significant effect for treatment 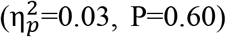, and a non-significant interaction effect 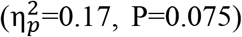. For relative peak torque (i.e., percent of isometric), there was a significant effect for velocity 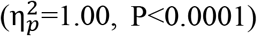, a non-significant effect for treatment 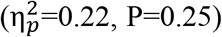, and a significant interaction effect 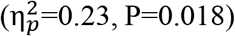. Based on post-hoc testing, relative peak torque was significantly higher (P=0.0004) in the NO_3_^−^ vs. the placebo trial at the highest velocity of 6.28 rad/s. Finally, for peak power there was a significant effect for velocity 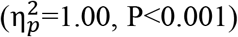, a non-significant effect for treatment 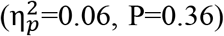, and a significant interaction effect 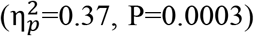. Post-hoc testing indicated that NO_3_^−^ ingestion significantly increased (P<0.0001) peak power at 6.28 rad/s.

**Table 3.** Effects of dietary NO_3_^−^ on peak knee extensor torque and power as a function of velocity in healthy 65-79 y old men and women.

|  |  | Angular velocity (rad/s) |  |  |  |  |
| --- | --- | --- | --- | --- | --- | --- |
|  |  | <u>0</u> | <u>1.57</u> | <u>3.14</u> | <u>4.71</u> | <u>6.28</u> |
| Peak torque<br>(Nm/kg) | Placebo | 1.94 | 1.42 | 1.02 | 0.75 | 0.60 |
|  |  | ±0.11 | ±0.09 | ±0.08 | ±0.07 | ±0.06 |
|  | Nitrate | 1.90 | 1.38 | 0.99 | 0.75 | 0.65 |
|  |  | ±0.11 | ±0.08 | ±0.07 | ±0.06 | ±0.06 |
| Peak torque<br>(% of isometric) | Placebo | 100.0 | 72.9 | 52.3 | 38.2 | 30.7 |
|  |  | ±0.0 | ±1.4 | ±2.2 | ±2.3 | ±2.0 |
|  | Nitrate | 100.0 | 73.1 | 53.2 | 40.4 | 34.5† |
|  |  | ±0.0 | ±2.1 | ±3.2 | ±3.4 | ±2.8 |
| Peak power<br>(W/kg) | Placebo | 0.00 | 2.04 | 3.28 | 3.77 | 3.50 |
|  |  | ±0.00 | ±0.13 | ±0.23 | ±0.31 | ±0.41 |
|  | Nitrate | 0.00 | 1.96 | 3.22 | 3.84 | 3.82‡ |
|  |  | ±0.00 | ±0.11 | ±0.20 | ±0.28 | ±0.37 |

Note. Values are mean±SE for n=12.
Nitrate trial significantly higher than placebo trial: †P<0.001; ‡P<0.0001.

Data for absolute and relative changes in Vmax and Pmax are shown in Figures 1 and 2, respectively. Dietary NO_3_^−^ ingestion increased Vmax from 10.1±0.6 to 11.2±0.6 rad/s 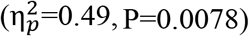, i.e., by 1.1±0.3 rad/s (Fig 2., *top panel*; 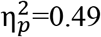, P=0.0079) or 10.9±3.5% (Fig. 2, *bottom panel*; 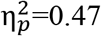, P=0.0099). Correspondingly, Pmax increased from 3.90±0.35 to 4.04±0.34 W/kg 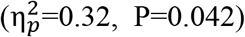, i.e., by 0.14±0.06 W/kg (Fig. 3, *top panel*; 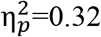, P=0.042) or 4.4±2.3% (Fig. 3, *bottom panel*; 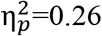, P=0.076).

**Figure 1.**
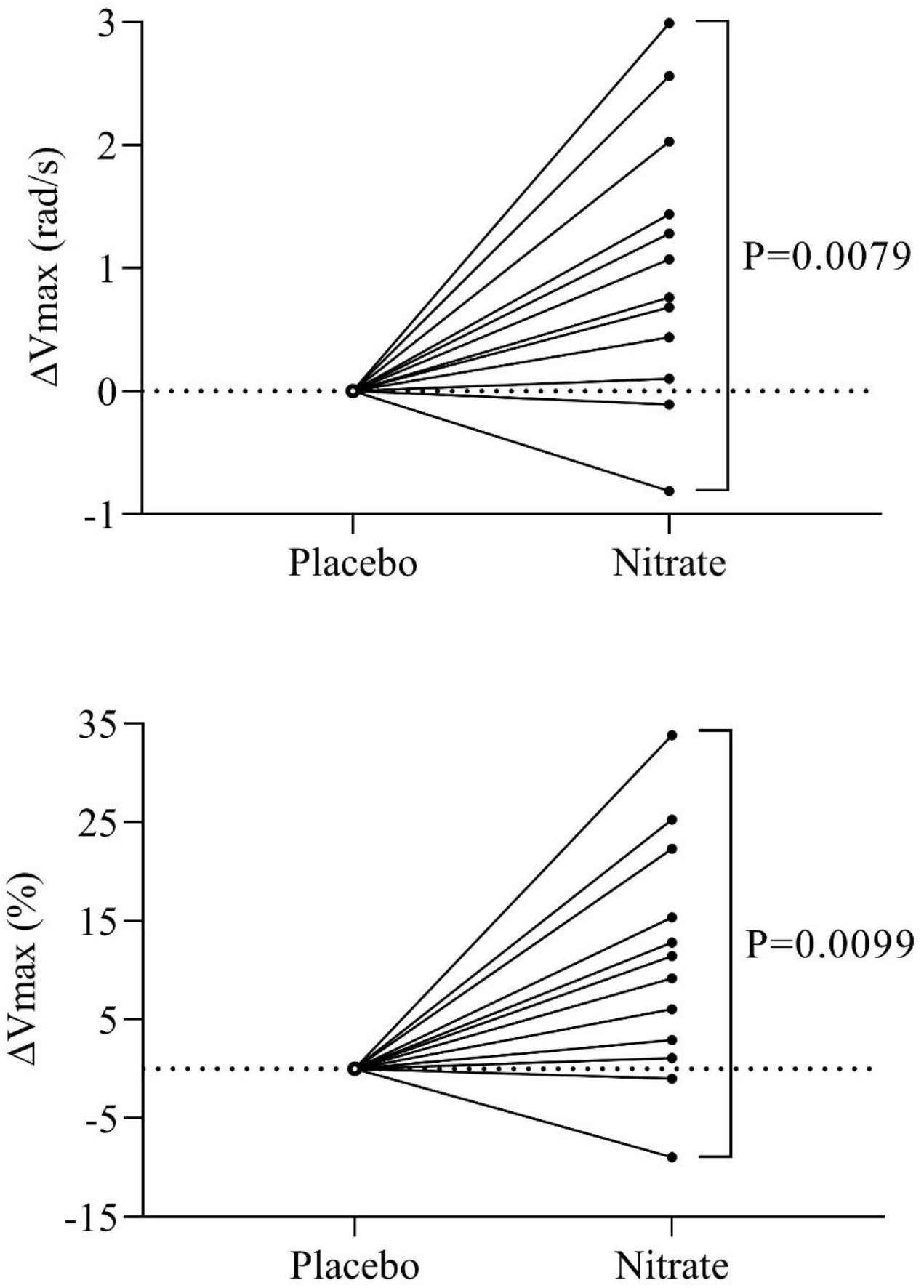
Absolute (*top panel*) and relative (*bottom panel*) changes in the maximal speed of knee extension (Vmax) in response to acute dietary NO_3_^−^ ingestion in healthy 65-79 y old men and women (n=12).

**Figure 2:**
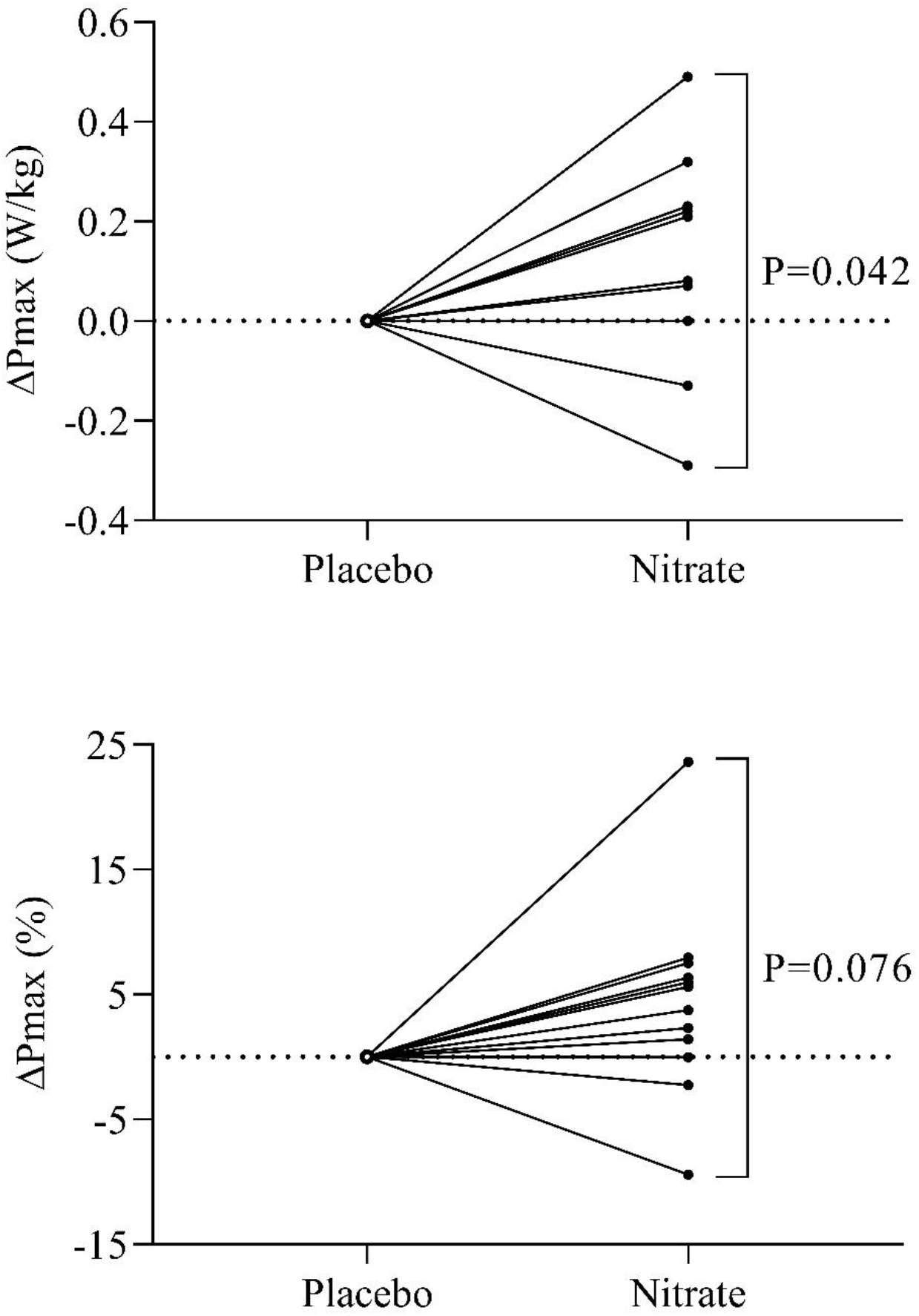
Absolute (*top panel*) and relative (*bottom panel*) changes in the maximal power of knee extension (Pmax) in response to acute dietary NO_3_^−^ ingestion in n=12 healthy 65-79 y old healthy men and women (n=12).

There were no significant differences between the placebo and NO_3_^−^ trials in peak power (3.49±0.30 vs. 3.51±0.31 W/kg; 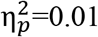, P=0.80), average power (1.08±0.08 vs. 1.10±0.09 W/kg; 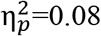, P=0.33), total work (37.0±2.8 vs. 37.6±3.2 J/kg; 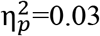, P=0.58), or fatigability (i.e., ratio of work during last 1/3^rd^ vs. first 1/3^rd^; 0.43±0.04 vs. 0.43±0.04; 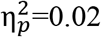, P=0.66) during the 50 contraction fatigue test.

## Discussion

The purpose of the present study was to test the hypothesis that acute dietary NO_3_^−^ supplementation would improve muscle speed and power in healthy older men and women. In keeping with our previous studies (21–24), we found that acute NO_3_^−^ ingestion increased Vmax and Pmax in older subjects by ~11 and ~4%, respectively. Although modest in size, the potential clinical significance of these changes is tremendous. Between the ages of 25 and 72 y, for example, the maximal velocity of unloaded knee extension has been reported to decrease by 27%, or by ~0.6% per year on average (35). Across a similar age range, peak knee extensor power was found to decrease by ~0.3% per year in men and ~0.6% per year in women (36). The dietary NO_3_^−^-induced increases in muscle speed and power observed in the present study are therefore functionally equivalent to acutely reversing the effects of approximately *one to three decades* of aging. Although greater attention is generally paid to age-related declines in muscle strength, age-related reductions in muscle speed have been found to play a greater role in explaining functional deficits in older mobility-limited individuals (37). Similarly, decreases in muscle power, i.e., the product of speed and force, with aging have been reported to be a more powerful predictor of functional decline than decreases in force (strength) alone (3). The significant improvements in Vmax and Pmax that we observed following acute NO_3_^−^ ingestion could thus have an enormous impact on the independence and quality of life of older individuals.

Although the dietary NO_3_^−^-induced in muscle function observed in the present are likely to be highly clinically significant, they are in fact similar in magnitude to those we have previously found for healthy young and middle-aged subjects (21,22). This observation runs counter to our hypothesis that older subjects would benefit *more* from dietary NO_3_^−^. This is presumably due to the fact that the healthy older men and women in the present study were not overtly NO deficient. Their baseline breath NO levels, for example, were comparable to those we have found previously in younger subjects (21,22). Their baseline plasma NO_3_^−^ and NO_2_^−^ concentrations were also similar to those we have measured previously (22). Additional studies will therefore be required to determine whether dietary NO_3_^−^ is even more effective at enhancing muscle contractile function in older men and women in whom NO bioavailability is more clearly reduced (and/or in those who are frail).

In contrast to the significant improvements observed during the force-velocity testing, acute NO_3_^−^ ingestion had no effect on peak power, average power, total work, or fatigability during the 50 contraction fatigue test. In keeping with this finding, Kelly et al. (19) previously reported that NO_3_^−^ supplementation does not alter muscle energetics during low or high intensity knee extensor exercise in healthy older men and women. Similarly, Siervo et al. (20) found that dietary NO_3_^−^ does not improve time-to-exhaustion during an incremental cycle ergometer exercise test in older subjects. More research into the effects of NO_3_^−^ supplementation on exercise capacity in this population is warranted. It is possible, for example, that older individuals may require a higher dose of NO_3_^−^ to elicit improvements in the capacity to perform sustained, high intensity exercise. Nonetheless, our data suggest that muscle contractile properties are more sensitive/responsive to NO_3_^−^ intake than resistance to fatigue during repeated contractions.

Along with the absence of changes in fatigue resistance during high intensity exercise, there were also no significant differences in blood pressure between the placebo and NO_3_^−^ trials. This observation differs from a number of previous studies (38,39), including some of older subjects (19,40). There is evidence, however, that aging may attenuate the vasodilatory effects of dietary NO_3_^−^ (41,42). Furthermore, there was a tendency for the *change* in systolic blood pressure to be greater after NO_3_^−^ ingestion (−9±2 vs. −3±2 mmHg; 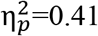, P=0.12), suggesting the lack of difference in absolute blood pressure may represent a type II error. Nonetheless, as with the fatigue test data discussed above our divergent results with respect to the impact of NO_3_^−^ intake on muscle contractile function and on blood pressure indicate that the former is more likely to show beneficial effects in older subjects.

There are a number of limitations to the present investigation. As indicated above, the small sample size may have precluded us from detecting statistically significant changes in blood pressure in response to dietary NO_3_^−^. Furthermore, although we recently hypothesized that dietary NO_3_^−^ enhances contractile function by altering Ca^2+^ release and/or sensitivity in muscle (43), the present study does not speak to this question. Along the same lines, while we have demonstrated that acute NO_3_^−^ intake is efficacious in enhancing muscle speed and power in older subjects, the optimal dose for improving this or other important physiological responses in this population remains to be identified. Further research will also be required to determine whether the acute responses that we have observed are maintained, or perhaps even magnified, by continued supplementation. Finally, it would be of interest to determine whether the effects of NO_3_^−^ supplementation are additive to, or perhaps even synergistic with, the effects of resistance exercise training.

In summary, we have demonstrated that acute ingestion of 13.4 mmol of NO_3_^−^, in the form of a concentrated BRJ supplement, significantly increases muscle speed and power in healthy, 65-79 y old men and women. This improvement in muscle contractility may help offset the declines in functional abilities and hence quality of life and independence that often accompany aging.

## Acknowledgments

This study was designed by A.R.C. and L.R.P.; experiments were performed/data were collected by all authors; data analysis, interpretation, and manuscript preparation were performed by A.R.C.

